# “Leap before you look”: Conditions that promote implicit visuomotor adaptation without explicit learning

**DOI:** 10.1101/2022.07.12.499675

**Authors:** Tejas Savalia, Rosemary A. Cowell, David E. Huber

## Abstract

When learning a novel visuomotor mapping (e.g., mirror writing), accuracy can improve quickly through explicit learning (e.g., move left to go right) but after considerable practice, implicit learning takes over, producing fast, natural movements. This implicit learning occurs automatically, but it has been unknown whether explicit learning is similarly obligatory. Using a reaching task with a 90-degree rotation between screen position and movement direction, we found that explicit learning could be “turned off” by introducing the rotation gradually (increments of 10-degrees) and instructing participants to move quickly. These specific conditions were crucial, because both explicit and implicit learning occurred if the rotation occurred suddenly, if participants were told to emphasize accuracy, or if visual feedback during movement was removed. We reached these conclusions by examining the time course of learning (e.g., whether there was fast improvement followed by a long tail of additional improvement), by examining the aftereffects of learning when the rotation was abruptly removed, and by using formal model comparison between a dual-state (explicit and implicit) versus a single-state learning model as applied to the data.

**Author summary:** In some situations, the relationship between motion direction and what we see is different than normal. For instance, try using a computer mouse that is held sideways (a 90-degree rotation). When first encountering this situation, people move carefully, using explicit strategies (e.g., move right to go up). However, after many learning trials, motion becomes automatic (implicit) and natural. Prior results found that implicit visuomotor learning always occurs with enough experience. In our study, we found that this is not true of explicit visuomotor learning; in some situations, explicit learning can be turned off. More specifically, we found that this occurs when the novel visuomotor situation is: 1) introduced gradually (e.g., a gradual introduction of 90-degree rotation in steps of 10 degrees); 2) when there is pressure to move quickly; and 3) with real-time onscreen views of the motion path. If any of these three components are missing, then people use explicit learning. These conclusions were reached by examining the time course of learning (e.g., whether there was both fast and slow learning as assessed with mathematical models) and by examining the tendency to automatically move in the opposite direction from the rotation when the rotation is abruptly removed after learning.

## Introduction

The spatial mapping between visual signals and motor actions is arbitrary and must be learned. For example, the retinal image is upside down and mirror reversed, and yet for a baby learning to interact with the world, this is no more or less intuitive than if the inversion and reflection were not present – retinal positions are arbitrary from the perspective of the brain, and through extensive experience we simply learn that to grasp an object appearing at the top right of the retinal image, we should move our hand downward and to the left. In some situations, the learned spatial mapping between vision and action is abruptly altered, requiring explicit correction. For instance, the first time that a new driver backs up a car using a rear-view mirror or camera, they need to purposely do the opposite of their natural tendencies. With practice, such novel mappings are implicitly learned, and action becomes automatic.

In laboratory studies, sensorimotor learning has been studied using paradigms that distort visual or motor feedback while participants perform reaching tasks, such as with kinematic distortion via force field manipulations [1–4], locomotor manipulations [5], visual distortions with prism glasses [6], or visuomotor adaptation tasks that involve center-out reaching under a rotated mouse-to-cursor mapping [7–19]. Results from these studies suggest there are two qualitatively different kinds of sensorimotor learning – one explicit and one implicit – that operate simultaneously, trading off with each other. The fast-to-learn, explicit system (i.e., a system that quickly adapts to the new sensorimotor mapping) produces slow but accurate responses early in learning. The slow-to-learn, implicit system guides fast automatic actions, but only after considerable practice. Studies supporting this distinction have, for instance, compared conditions that give aiming instructions about the sensorimotor perturbation [9,20], compared conditions with versus without online visual feedback [21], or compared gradual versus abrupt introduction of a sensorimotor perturbation [7]. Aftereffects of a sensorimotor perturbation are often examined by abruptly returning the sensorimotor mapping to the standard situation in a “washout” phase [7], or by examining whether sensorimotor learning transfers to other actions [22].

A particularly clear demonstration of the explicit/implicit learning distinction is found in the study of Mazzoni and Krakauer [20], in which participants were explicitly told to aim 45 degrees counterclockwise to compensate for a 45-degree clockwise visuomotor rotation in the mapping between visually displayed targets and reaching direction in a pointing task. This instruction allowed almost perfect performance soon after rotation, but, paradoxically, movements slowly drifted in the direction of “overcorrection” (e.g., even further counterclockwise than was needed to counteract the rotation). In other words, the implicit system continued to slowly adjust to the rotation even though the explicit strategy was sufficient for good performance. Thus, even when explicit learning is maximized through an explicit aiming strategy, implicit learning still occurs even if implicit learning results in worse performance. In other words, implicit motor adaptation is automatic. In the current study we ascertain whether the complement is true: Does explicit learning always occur, or can it be turned off in the right circumstances?

As applied to learning a novel visuomotor rotation, “dual state” models of motor learning assume that actions on each trial reflect the summation of the rotation estimates provided by two different learning systems (aka, two “states” that estimate the required rotation) [3,9,23]. When referring to the dual state model we adopt the terminology of the “fast-learning system” for explicit learning and the “slow-learning system” for implicit learning [23]. A point of potential confusion, which we clarify here, is that the fast-*learning* system results in more slowly executed movements, on any given trial (for example, it may dominate when participants are forced to wait before acting [26]), whereas the *slow-learning* system produces knowledge or representations that are deployed more quickly, i.e., allowing faster movements. The model assumes that both systems operate in the same manner, differing only in how quickly each system learns and forgets the novel rotation. In other words, the model is a descriptive account of what happens when a fast-learning and slow-learning system compete with each other to produce behavior. Because the rotation estimates from each system are summed, accurate motion could reflect a perfect estimate of the rotation in the fast system combined with no rotation in the slow system, or vice versa, or a situation in which each system provides a partial estimate of rotation, with these adding up to the correct total. One consequence of this additive assumption is that learning curves exhibit a quick reduction in error as the fast system rapidly develops an accurate estimate of rotation, followed by a longer tail as the estimate of rotation gradually shifts from the fast to the slow system [14]. Because the slow system not only learns slowly but also forgets slowly, the dual state model successfully explains the slow return to accurate performance upon abrupt removal of the rotation, spontaneous recovery as well as savings during re-adaptation [3,24,25]. Supporting the assumption that the fast system reflects explicit learning or strategies, the fast system has been found to reflect the explicit aim reported by participants [14] and to capture conditions where participants are given a strategy to counter the perturbation [9].

Most studies supporting the dual state model instruct participants to be both fast and accurate, rather than emphasizing only speed. Because there is always some emphasis on accuracy, this may lead participants to use some form of explicit aiming strategy. Thus, these instructions may be critical for ensuring that the explicit system plays some role in adapting to a novel sensorimotor perturbation. In other words, the success of the dual state model may reflect the combination of obligatory implicit learning [9,20], which is captured by the slow-learning state [23], and an explicit aiming strategy in light of instructions that emphasize accuracy, which is captured by the fast-learning state. Providing support for this claim, forcing participants to wait before moving has been found to increase accuracy through explicit aiming strategies [26]. Conversely, because the fast- and slow-learning systems interact to produce a combined error signal in the dual state model [27], pressure to respond quickly should lead to an increased contribution of the slow-to-learn (but fast to deploy) implicit system [15]. To ascertain whether an emphasis on speed might reduce or even eliminate the contribution from the explicit fast system, the current study examined visuomotor adaptation to a 90° rotation in separate conditions that emphasized either fast motion or accurate motion.

Another important factor for the explicit system may be whether errors are sufficiently large as to be noticeable. For instance, if a large rotation is abruptly introduced, participants will become explicitly aware of the rotation, and this awareness may underlie learning in the explicit system even in the absence of pressure to be accurate. One way to address this possibility is by introducing the rotation gradually in increments such that participants are never explicitly aware of any large discrepancies between their movements and the desired outcome (e.g., the gradual introduction of a 90° counter-clockwise rotation in steps of 10°, with time to implicitly adapt to each additional change of 10°). Prior work with a gradual introduction of rotation found that it results in a larger washout aftereffect [7,28]. That is, when participants adapt to a 90° visuomotor rotation that is introduced all-at-once (sudden perturbation) participants tend to make fewer errors when the rotation is subsequently removed as compared to when the 90° rotation in introduced in small increments (gradual perturbation). These results are consistent with the claim that a gradual introduction of rotation more effectively promotes implicit learning, which is then more slowly forgotten, producing greater errors upon removal of the rotation. This claim finds additional support in the finding that that Parkinson’s patients can successfully adapt to a gradual perturbation but not a sudden perturbation [29]. In other words, lacking effective explicit movement control, Parkinson’s patients struggle to adapt to a sudden introduction of a large rotation, but their intact implicit motor control system can nonetheless adapt gradually, if environmental conditions permit.

Beyond an emphasis on speed and the introduction of the perturbation gradually, another factor that may enhance implicit adaptation is continuous onscreen cursor feedback during motions as compared to feedback that shows only the final result of the motion [10,14]. In other words, allowing participants to see only the end result encourages the use of explicit aiming strategies. To maximize the contribution of the implicit system, our first study always presented continuous cursor feedback during each motion (see *Methods*).

In our first study, we ask whether visuomotor adaptation necessarily involves the fast-to-learn explicit system. We used a 2×2 between-subjects design in a center-out reaching task under visuomotor rotation to determine how speed versus accuracy emphasis changes the course of learning. We also asked whether the speed/accuracy manipulation interacts with a manipulation of rotation, namely, whether the rotation is introduced gradually versus suddenly. In particular, the combination of speed emphasis and gradual introduction of the rotation may eliminate use of explicit learning. This combination may encourage participants to begin moving (i.e., “leap”) without explicit correction for the motion direction. That is, rather than “look before you leap”, this may produce “leap before you look”. We examined the time course of learning to assess whether there was, or was not, an initial rapid correction, such as expected with explicit learning. We also examined behavior when the rotation was abruptly removed after extensive training. This abrupt removal is likely to induce use of the explicit system in all conditions, and the question of interest was how this rapid unlearning of the rotation compares to the initial speed of learning the rotation, and whether some conditions produce a greater aftereffect in this washout phase. To ascertain whether the fast-learning system is needed to explain performance, we applied both the dual state model and a single state model to the initial learning results, using formal model comparison to ask whether both the fast- and slow-learning systems are necessary [3,30] in each of the four experimental conditions. To preview our results, we found that the fast-to-learn explicit system is not always engaged. Our conclusion that the explicit system is optional complements prior investigations of the implicit system, which concluded that the implicit system is obligatory [9,20].

## Experiment 1

### Methods

#### Participants

Sixty undergraduate students from the University of Massachusetts, Amherst participated in the study. Participants received course extra credit as compensation for time spent in the study. There were 15 participants in each of the four between-subjects conditions. The study was approved by the Institutional Review Board at University of Massachusetts, Amherst. All participants gave informed consent prior to the start of the experiment.

#### Paradigm

Each participant was seated in front of an LCD screen that displayed experimental stimuli. A trackpad with an active area of 10×6.25 inches was placed in front of the monitor and motor responses were made with the preferred hand using a stylus on the trackpad. Movements were recorded using Psychtoolbox routines within MATLAB, run on a Windows PC.

The experiment design is illustrated in Figure 1. On each trial, participants used the stylus on the trackpad to move the onscreen cursor from its center starting position to the displayed target. Each trial started with presentation of a small circle (“cursor”) at the center of the screen. Next, a target circle appeared 0.75 to 1.25 seconds later (stimulus onset asynchrony drawn from a uniform distribution). The target appeared at one of the four possible locations, which were the four diagonal directions relative to the central cursor (upper-left, upper-right, lower-left, and lower-right). Participants moved the cursor to the target in order to complete a trial. They were provided with continuous feedback of cursor movements, which allowed them to correct their motion until they hit the target. If the participant lifted the stylus mid-trial (unsuccessful movement) or exceeded the active area of the trackpad, the cursor reset to the center. Only successful movement trajectories from the center to the target were used for further analyses. Once the target was reached, the cursor reset to the center of the screen, indicating the start of the next trial. The first point of contact of the stylus to the trackpad was mapped to the cursor position at the center of the screen. All subsequent on-screen cursor positions were displayed relative to this first point of contact. This allowed participants to start a movement from any point on the trackpad without having to find the trackpad center at the start of each movement.

**Fig 1.**
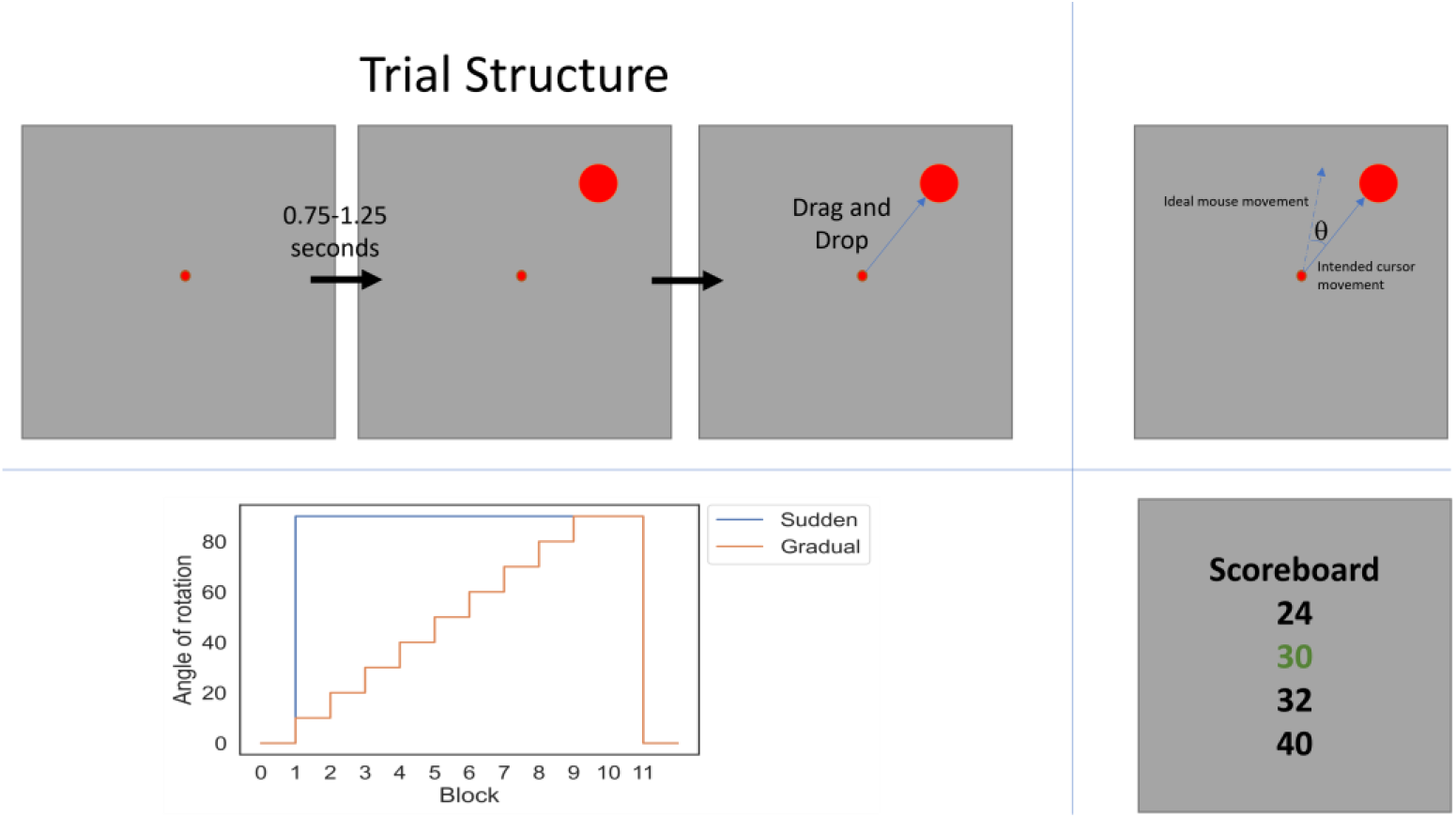
*Top Left:* Trial sequence for an example trial with an upper-right target: Each trial began with a red circle in the center of the screen, which was the starting point for movements. After a variable delay of one second on average, a target circle appeared at one of four predefined locations on the screen (in one of four diagonal positions relative to the central dot: upper-right, upper-left, lower-left, or lower-right). Using a stylus, participants dragged the cursor from the center of the screen to the large target circle in a drag and drop motion. *Top Right:* Visuomotor perturbation. The stylus-cursor mapping was perturbed such that the on-screen cursor direction was rotated in the clockwise direction by angle (*θ*) from the direction of stylus movement (the ideal movement direction for an error-free movement was thus rotated counterclockwise). *Bottom Left:* Sudden/gradual manipulation: The sudden perturbation schedule presented 10 blocks of 90° rotation, whereas the gradual perturbation schedule gradually achieved a 90° rotation by implementing 10° increments across 9 blocks with the final two blocks inducing 90° rotations. Each block included 64 trials. *Bottom Right:* Speed/accuracy manipulation: The participant’s score, shown in green, was calculated based on either speed or accuracy for all trials within a block. This feedback appeared in a scoreboard shown at the end of each block. The other scores in the scoreboard were fictitious and were randomly drawn from a normal distribution centered around the participant’s score.

The experimental paradigm consisted of 12 blocks of 64 trials each, divided into 3 unequally sized phases (1 block, 10 blocks,1 block). Each block of 64 trials consisted of 16 reaching movements to each of the four targets. Trials in a block were pseudo-randomly ordered with four sub-sequences of 16 trials, which each contained a random order of 4 reach movements to each of the 4 targets. After a short set of familiarization trials for which the experimenter was present to ensure successful understanding of the task, the first phase, “baseline”, lasted for one block of 64 trials. During the baseline phase, the mapping between trackpad and onscreen directions was standard (un-rotated). The second phase, “rotation”, lasted for 10 blocks of 64 trials. The mapping was distorted by rotating the on-screen cursor by a maximum of 90° clockwise relative to the position of the stylus on the trackpad. The final phase, “washout”, lasted for one block of 64 trials without rotation (identical to the baseline phase). The start of each block was self-paced – the participant indicated by stylus click when they were ready to start a new block.

Participants were randomly divided into 4 groups of 15. Two of the groups were asked to make their movements as quickly as possible while the other two were asked to make their movements as accurately as possible. To reinforce the speed/ accuracy instruction, at the end of each block, participants were shown a scoreboard indicating their performance in that block relative to four other scores of hypothetical competitors. Block scores were calculated by adding up a participant’s performance across all 64 trials within a block. For speed emphasis participants, the score for each trial was calculated as the inverse of the time between the target’s appearance and the time at which the target was reached. For accuracy emphasis participants, the score for each trial was the inverse of Euclidean distance in pixels between each sample of the participant’s trajectory and the closest point on a straight line from the center to target. In the scoreboard shown at the end of the block, the participant’s total score (indicated by green text) was always shown at positions 2-4. The other three scores were created by taking random draws from a normal distribution that was centered on the participant’s score. If a participant’s score happened to be the highest in the random draw of 4 scores, scores were resampled till there was at least one random score better than the participant’s, thus giving the participant some incentive for continued improvement.

All 60 participants adapted to a 90° clockwise rotation. In the rotation phase, one of the speed groups of participants and one of the accuracy groups of participant were assigned to receive a “Sudden” rotation. These groups experienced the 90° rotation for all 10 blocks of the rotation phase. The other two groups were assigned to receive a “Gradual” rotation by gradually building up to the to the 90° rotation in nine increments of 10° across nine blocks, with the tenth block holding steady at 90°.

#### Movement Analyses

Cursor trajectories and movement times were recorded for each trial. Movement initiation time was calculated as the time between the appearance of the target and the time it took for the cursor to cross 5% of the Euclidean distance to the target. Movement time was calculated as the time between the point when 5% distance was covered and the time when the target was reached.

Angular error was calculated as the average absolute angular error for each sampled point along the motion path. For each sampled data point, the absolute angle was calculated between a vector from that point and the next sampled point and a vector from that point to the target (i.e., the ideal direction). To reduce noise in the data, all further statistical analyses (but not model analyses) were performed on a smoothed version of angular error. Smoothing was calculated using a weighted average of angular error across trials with weights determined by a Gaussian kernel with standard deviation of 2 trials.

#### Model Fitting

Both the single state and dual state models were fit to the data from the rotation phase, separately for each participant, to determine which participants required both learning systems for an adequate description of their learning. The baseline block was not included in model fitting because performance was highly accurate and there was nothing learn (there was no rotation, and the model estimate of rotation was initialized to zero). The washout block was not included because it would bias model comparison in favor of the dual state model considering that all participants were likely to become explicitly aware of the sudden removal of rotation. Critically, these fits were not to the absolute angular error but rather to the signed angular error, thus capturing both increases and decreases in rotation estimates based on the trial-by-trial behavior (e.g., if a model overcorrects for the rotation, which corresponds to an estimated rotation that is larger than it should be, this negative error signal will reduce the rotation estimate). To fit data, the model took on parameter values that were constant across the entire trial sequence of the study (considering that the fitted parameters comprised a retention rate and learning rate for the model’s estimate of rotation, this of course allowed the rotation estimate itself to change across trials). The model produced a sequence of predicted angular errors with these errors arising from the actual trial sequence (e.g., in the experimental design, the sequence of actual rotation values). We explain first the single state model considering that the dual state model is an extension of the single state model. For the single state model, error experienced on any given trial is the difference between the rotation estimate based on the experience in the previous trial and the actual rotation as dictated by the experimental design sequence for that trial (Eq. 1). The first trial starts with an estimate of no rotation. The equations for this model dictate that the estimate of rotation, *r_est_*, at the end of the current trial, *t*, is a weighted average (parameters *A* and *B* determining the weights) between the previous estimate of the rotation and the currently experienced error (Eq. 2); the currently experienced error (Eq. 2). [3,31].

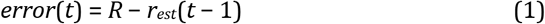

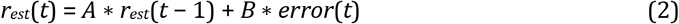

Here *R* is the rotation for the current trial as dictated by the experimental design, *r_est_*(*t*) is the estimate of rotation on trial *t*, the parameter *A* represents a “retention” factor and the parameter *B* represents a learning rate.

We also fit the dual state model to error data [3]. The dual state model postulates that there are two interactive systems involved in successful adaptation to a rotated stylus-cursor mapping [3,9,19,23] as described by the following equations.

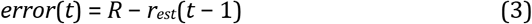

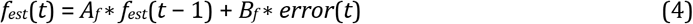

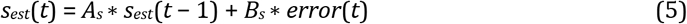

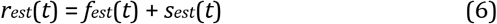

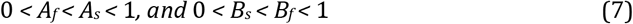

The total estimate of rotation is modeled as the sum of two independent systems: A “fast” system (*f*) that learns quickly and forgets quickly and a “slow” system (*s*) that learns slowly but retains the learned mapping for longer than the fast system.

Data fitting was done in two stages, first using all 640 trials from the rotation phase, with three separate model comparison measures of the results (chi-square, AIC, and BIC), followed by cross-validation that randomly divided the data into training and test sets. To calculate the likelihood of each observed error for the rotation phase, it was assumed that the predicted error calculated in equations 1 and 3 reflected the mean of normally distributed error, with a free parameter for error variance. In this way, to the extent that the model was able to capture the observed data, the variance could be set to smaller values, thus increasing the likelihood of the observed data. In the first stage, which fit all trials from rotation learning, a global best-fit for each participant was found by running multiples instances of simplex minimization, with each instance using a different set of starting parameters, determined by a small grid of parameter values. This was done separately for each model and the best fitting parameters values were used to generate three different model comparison measures (chi-squared, AIC, and BIC). In the second stage, which used cross-validation, the trials for each participant were divided into a training set of 90% (576) trials picked at random with the remaining 10% (64) trials serving as the test set. 100 different train-test sets were created for each participant and fit separately using the simplex minimization routine. For each cross-validation fit of the training data, the parameter values from the first stage were used as the starting point, thus ensuring that the parameter values were “in the ballpark” for each sample of training data. Model performance for the second stage was assessed by examining how well each model predicted, a priori, the 100 different held-out samples of test data for each participant, using the best-fit parameters from the corresponding training samples. These predictions for the held out data were not assessed using point estimates of the angular errors but rather the likelihood of the observed held-out data according to the model with parameters that had been fit to the training data. In this way, the models were also assessed in terms of their ability to capture the reliability of the predictions (e.g., whether the error variance parameter was well-matched to variability in the predicted data).

### Results

#### Performance after Learning

The first 9 blocks of rotation learning were non-equivalent between the sudden and gradual and sudden groups, because each of the 9 blocks introduced an additional rotation change for the gradual condition but not the sudden condition. This non-equivalence made it difficult to compare the conditions during the first 9 blocks of rotation learning, although, as reported below, the time course of learning can be compared by applying learning models to the data. To compare conditions after learning, we analyzed the 10th block (final block of rotation) and 11th block (“washout”), which were identical across conditions. The 10th block was a continuation of the 90° rotation for all groups and the washout block was the sudden removal of the rotation for all groups. These analyses used a between-subjects 2X2 ANOVA with the factors of type of rotation learning (sudden/gradual) and performance emphasis (speed/accuracy), with either average absolute angular error or median latency as the dependent measure.

As shown in the left panel of Figure 2, for the angular error during the final block of rotation learning, there was a main effect of performance emphasis (*F* = 18.07,*p* < 0.001), with lower error for the accuracy emphasis groups, demonstrating that accuracy emphasis instructions produced more accurate movements. There was no main effect for the type of learning, i.e., sudden versus gradual (*F* = 1.54,*p* = 0.22), and no interaction between the two factors of performance emphasis and learning type (*F* = 0.92,*p* = 0.32). As shown in the right panel of the figure, the subsequent washout block produced very different results, with no apparent effect of performance emphasis (F = 2.75, p = 0.102) and yet there was a main effect for type of rotation (*F* = 13.39,*p* < 0.001), with greater error for the gradual rotation groups. This effect of gradual versus sudden rotation learning during the washout block replicates prior work [7]. These two factors did not interact during the washout block (*F* = 0.017,*p* = 0.89). Demonstrating that the elimination of the performance emphasis effect and the emergence of the type of rotation effect between blocks 10 and 11 were both reliable, a three-way ANOVA that also included block (final rotation, 10, versus washout, 11), revealed a significant interaction between block and performance emphasis (F = 4.8, p = 0.03) and a significant interaction between block and type of rotation (F = 10.89, p = 0.001).

**Fig 2.**
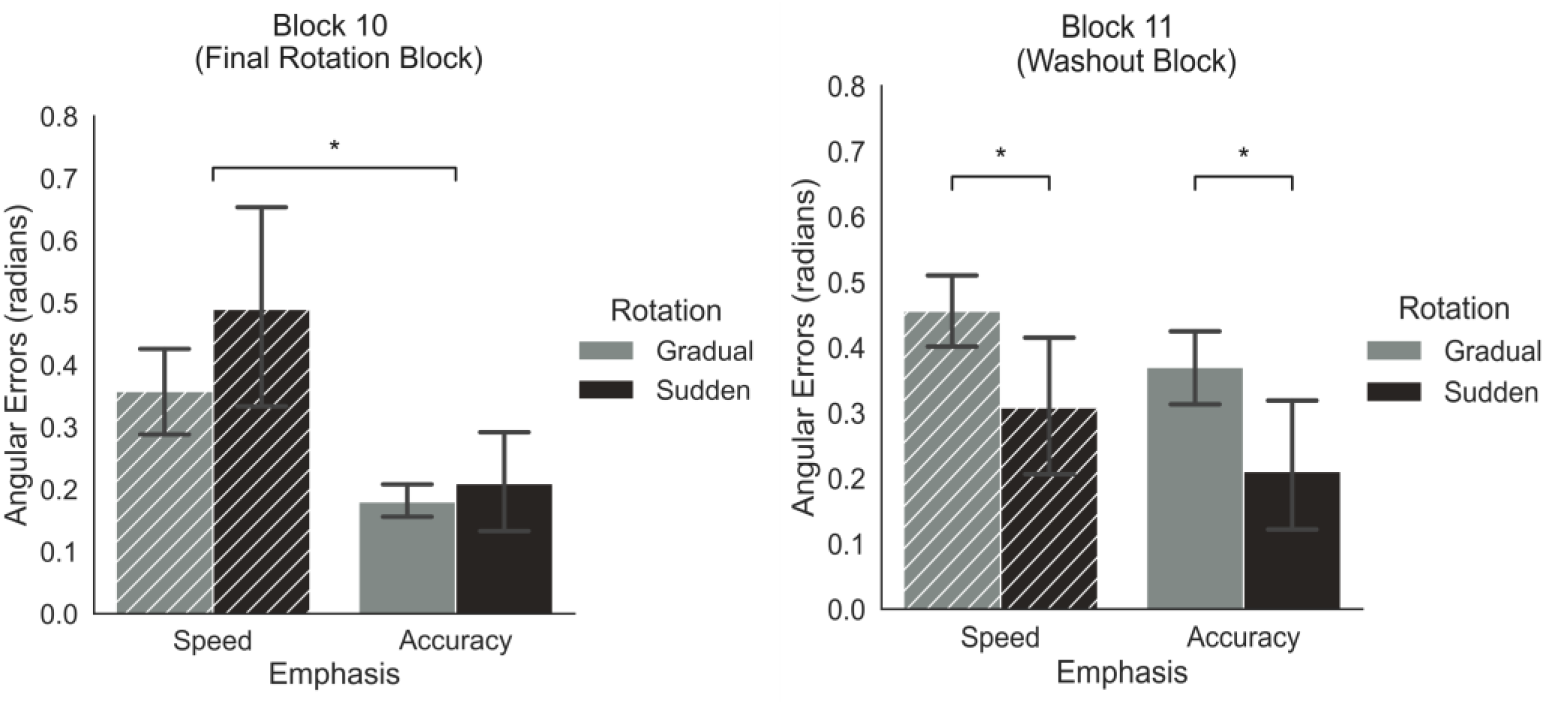
Accuracy at the end of learning in Experiment 1. *Left Panel* During the final block of rotation learning, participants in the speed conditions made larger errors than participants in the accuracy conditions for both the gradual and sudden introduction of rotation, demonstrating that the speed/accuracy instructions affected behavior as expected. *Right Panel* During the washout block, in which the rotation was removed, participants who had learned the rotation gradually produced larger errors than those who had learned the rotation suddenly.

To examine whether participants adhered to the speed/accuracy instructions, median latency was examined separately for the time to initiate movement and movement time. Although median initiation times were not affected by performance emphasis instructions (*t*_(58)_ = −0.85,*p* = 0.39), median movement times were affected by emphasis, with faster movements for the speed emphasis groups of participants (*t*_(58)_ =−2.22,*p* = 0.03). Thus, in conjunction with the accuracy results reported above, there was a speed-accuracy tradeoff when comparing the speed emphasis groups to the accuracy emphasis groups.

In summary, accuracy versus speed emphasis affected accuracy at the end of rotation learning and yet, in the washout block, the main finding was greater errors following the gradual introduction of the rotation, regardless of speed/accuracy emphasis. This pattern implies that a longer-lasting form of learning took place during the 9 blocks of rotation learning for the gradual introduction of the rotation. Next we turn to analyses of the time course of learning, using the single and dual state learning models to determine whether this longer-lasting form of learning reflected the slow-to-learn implicit system and whether the fast-to-learn explicit system was needed in all conditions.

#### Learning Curves: Are Two Learning Systems Needed?

Our central question was whether both the explicit fast-to-learn system and the implicit slow-to-learn system are necessary or whether in some cases behavior is better explained using only the slow-to-learn implicit system. To address this question we used formal model comparison, comparing the dual state model to the single state model as applied to the trial-by-trial data from the 10 blocks of rotation learning, as seen in Figure 3. Although the slow-to-learn implicit system may be an obligatory component of motor learning, as indicated by prior studies [10,14], the fast-to-learn explicit system may not be obligatory; situations that favor the single state model are those most likely to obviate the need for the explicit system.

**Fig 3.**
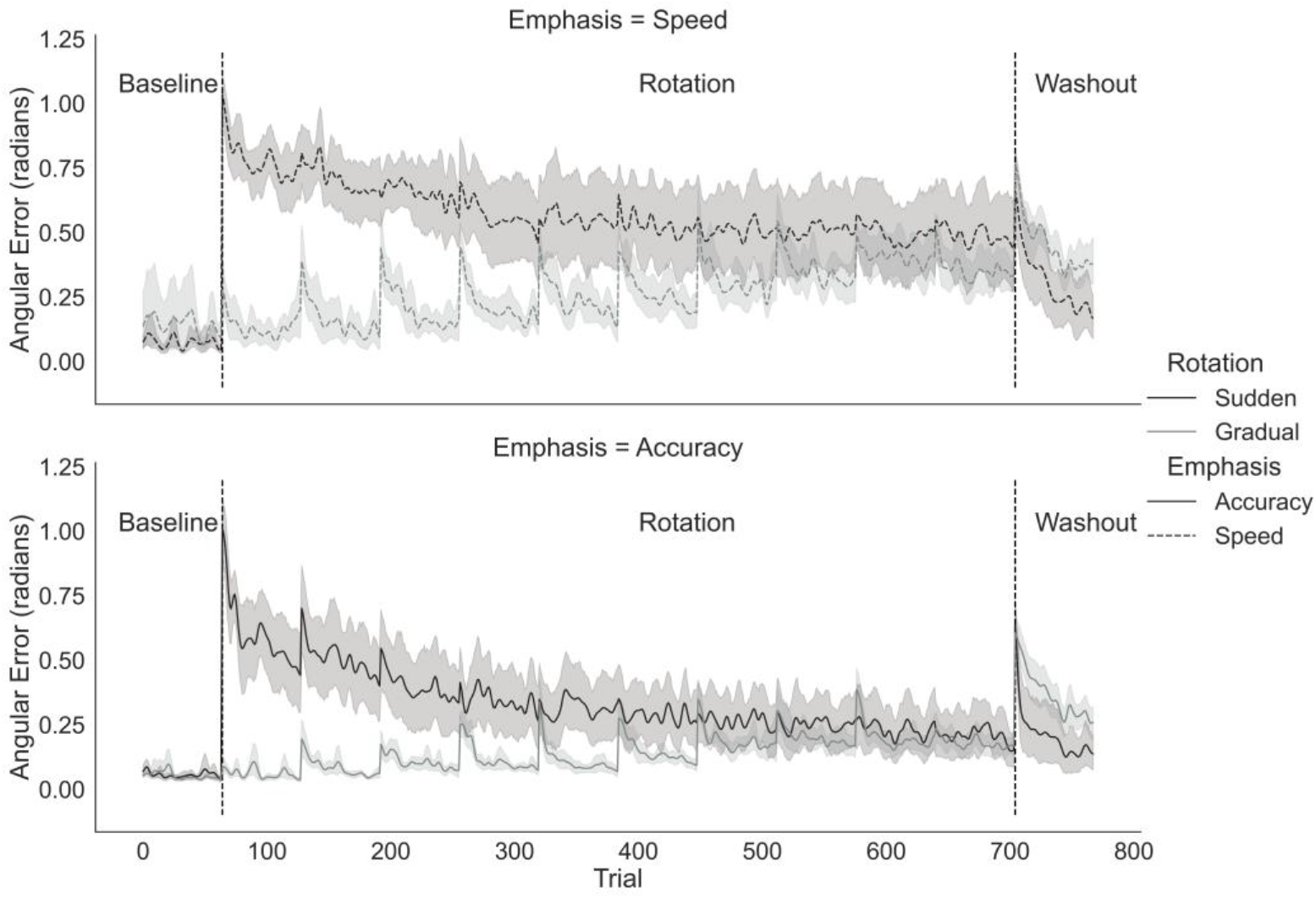
Average absolute angular motion direction error for the four groups of participants as a function of trial in Experiment 1. Darker lines indicate the groups that experienced a sudden introduction of the 90° rotation, and lighter lines the groups that were introduced to the rotation gradually. *Top Panel:* Results for speed emphasis groups (dotted lines). *Bottom panel:* Results for accuracy emphasis groups (solid lines). Shaded regions indicate the 95% confidence interval

The main difference between the models is that the dual state model can capture a learning curve that shows both very rapid improvements over just a few trials as well as much slower improvements over the course of hundreds of trials. Such patterns are readily apparent in the average results shown in Figure 3, for the sudden rotation conditions (black lines). In these conditions, after the 90° rotation is suddenly introduced at the 65th trial, the average angular error is initially close to 1 radian (or 57 degrees), but then this error is rapidly reduced to approximately .6 radians (accuracy emphasis) or .8 radians (speed emphasis) over the first 5-10 trials of Rotation learning. After this initial error reduction, a very slow time course of additional error reduction ensues. Something similar is seen in the gradual accuracy condition (solid grey line, lower plot), not in the first 5-10 trials of Rotation, but rather in the first 5-10 trials of every new block (every set of 64 trials) of Rotation learning, whenever a 10° rotational increment is introduced. It can perhaps be observed that, in the gradual speed condition (dotted grey line, upper plot), the period of error reduction in each block is more drawn out (i.e., the error reduction happens less sharply) than in the gradual accuracy condition (solid grey line, lower plot); this may provide an important clue to the involvement of the two systems in the different conditions. However, these descriptions are speculative and are based on average results, which can be misleading in light of different learning patterns for different participants (e.g., perhaps some participants learn quickly and others learn slowly, in which case the average pattern would show an initial rapid decrease followed by a long tail of additional learning as an artifact of averaging).

To test the nature of the learning curves, both models (dual state and single state) were fit separately to the learning curves of each participant to determine, for each participant, whether learning entailed both fast and slow learning (i.e., the dual state model), or whether learning entailed only slow learning (i.e., the single state model). The dual state model has more free parameters that can be fine-tuned to fit data, allowing it to capture a wider range of behaviors. Thus, a head-to-head comparison between these two learning models requires some sort of penalization for the extra flexibility of the dual state model [32]. Table 1 shows the results using four different penalization techniques. The values in Table 1 are the number (and proportion) of participants in each group whose behavior was better explained by the dual state model; the remainder were better fit by the single state model. For instance, where the table shows 0, this means that none of the participants in the corresponding group produced learning curves that were better explained by the dual state model. Cutting straight to the main result, all four methods of model comparison indicated that the dual state model was not needed for the specific combination of speed emphasis instructions and a gradual introduction of the rotation (i.e., for the “gradual speed” condition, the single state model provided a sufficient explanation of behavior without requiring a second state).

**Table 1.** Model Comparison Results for Experiment 1 Showing How Often the Dual State Model was Preferred

| Comparison<br>Metric | Sudden<br>Speed | Sudden<br>Accuracy | Gradual<br>Speed | Gradual<br>Accuracy |
| --- | --- | --- | --- | --- |
| Chi2 | 6 (40%) | 12 (80%) | 0 (0%) | 9 (60%) |
| AIC | 7 (46%) | 13 (87%) | 0 (0%) | 9 (60%) |
| BIC | 5 (33%) | 11 (73%) | 0 (0%) | 6 (40%) |
| CrossVal | 9 (60%) | 13 (87%) | 3 (20%) | 10 (67%) |

#### Different Methods of Model Comparison

First, we compared the two models using chi-squared tests. Because the single state model is nested under the dual state model, one can ask whether the single state model provides a significantly worse fit considering that it is a special case of the dual state model that removes two of the parameters. More precisely, there are two circumstances in which the dual state model is the same as the single state model: 1) if the slow-learning state learning parameter (Bs) is fixed at 0 (no learning) and hence the fast learning state remains; or 2) if the fast- and slow-learning states have the same rates of retention as the single state model (Af = As = A), and the learning rates of the slow- and fast-learning states add up to the learning rate of the single state model (Bf + Bs = B). In first case, the contribution of the slow-learning state is turned off (i.e., it always predicts a zero rotation), and the fast-learning state is the only state that learns the rotation.

However, we emphasize that in this first case, the constraint that 0 *< A_f_ < A_s_ <* 1,and 0 *< B_s_ < B_f_ <* 1 (Eq. 7) mandates that any learning loads onto the fast-learning state parameters, Af and Bf. Therefore, such a result emerging in the dual state model could in fact reflect that only the *slow-learning* state is in operation, with its parameters labelled as “fast-learning state” owing to model constraints. In the second case, the two states together learn (and forget) at the same speed as the single state model, and their individual estimates of rotation add up to be same value as that of the single state model.

For nested models, one test of whether the extra parameters of the more complicated model provide some benefit is based on the likelihood ratio test and the corresponding G-squared goodness of fit metric [33]; accordingly, we used a chi-squared test with two degrees of freedom to assess whether the single state model provided a significantly worse fit, separately for each participant. The top row of Table 1 shows the results of this nested model comparison, revealing that the extra free parameters of the dual state model were not warranted for any participant in the gradual speed group. In contrast, nearly all of the participants (80%) in the sudden accuracy group produced learning curves that were better explained by the dual state model. The sudden speed and gradual accuracy groups were more mixed, with a more moderate proportion of participants that were better explained by the dual state model than the single state model.

The chi-squared test for nested models assumes that the parameters can take on any value and yet this is not strictly true with the dual state model, because it imposes rank order constraints between the two states. For instance, it is not allowed to have one state with slower forgetting (A) but faster learning (B) as compared to the other state (the slow-learning state is constrained to always have both slower forgetting and slower learning, Eq. 7). Thus, nested model comparison somewhat unfairly handicaps the dual state model, which is not truly free to set its parameters to any value. There are a wide variety of techniques that can be applied to non-nested models and Table 1 also reports model comparison using the Akaike Information Criterion (AIC [34]), and the Bayesian Information Criterion (BIC [35]). It is commonly understood that AIC, which is based on predicting data, tends to favor more flexible models whereas BIC, which is based on identifying the more likely model, tends to favor less flexible models. Correspondingly, as seen in Table 1, the dual state model did somewhat better on average when using AIC and did somewhat worse on average when using BIC, as compared to the chi-squared results. However, for both AIC and BIC, it was still the case that no participant in the gradual speed condition produced behavior that was better explained by the dual state model.

AIC and BIC are parameter-counting measures, and assume that each additional free parameter entails the same degree of extra flexibility. However, some parameters are likely to be more important than others (e.g., inclusion of a second state might affect goodness of fit only for trials occurring early in learning, whereas the error variance parameter affects all trials). Furthermore, as mentioned above, the dual-state model may be overly penalized on the basis of a simple parameter count, because the rank order constraints between the two states (Eq. 7) reduce the extra flexibility provided by the second state. To seek a fairer model comparison metric, we examined available techniques for non-nested models that avoid parameter-counting (for a review see [32]), and from among these we opted to use cross-validation [36]), which is relatively assumption free and easy to implement through non-parametric sampling of the data. When using cross-validation, an overly flexible model will fit noise in the training data and produce a worse prediction for held out data. As with the chi-squared test, we asked whether the extra flexibility of the dual state was warranted. This framing of the question implies a default selection of the simpler model in the case of non-diagnostic data ([37]). Thus, the dual state model was selected to be the winner for a particular participant if it performed better than the single state model significantly more often, which corresponded to 59 of the 100 cross-validation predictions (Binomial test cut-off for p = 0.05, n = 100) when comparing the likelihood of the held out test data based on parameters fit to the training data. As seen in the bottom row of Table 1, the dual state model did somewhat better on average for cross-validation, as compared to the other methods of model comparison. However, only 20% of the participants in the gradual speed group were better fit by the dual state model. Thus, even when using a technique that accurately addresses the flexibility of the models, the addition of the fast-to-learn explicit system was not often needed in the gradual speed condition.

#### Is the Single State Implicit Learning?

In reaching conclusions from the model comparison results, we assumed that when a second learning state was not needed (i.e., when the dual state model failed to fit better than the single state model), the single state reflected the slow-to-learn implicit system. This interpretation was based on prior evidence that the implicit system is an obligatory component of learning [10,20]. Nevertheless, this assumption can be checked directly by examining the best-fitting learning and retention rate parameter values. If only one learning state was needed, did this single state correspond to a relatively slow time course of improvement? The time course of improvement for a state reflects both the learning parameter and the retention parameter. For instance, a high learning parameter would learn on every trial, but if the retention parameter were low, then the learning from the previous trial would be immediately forgotten, and there would be little improvement. The rate of accuracy improvement is related to the multiplication of the learning and retention parameters (e.g., it can be algebraically proven that this relationship is directly proportional for the change from the second to the third step of learning), and so we used this multiplication term in our analysis, as a proxy for rate of improvement. The results are shown in Figure 4, which plots side-by-side the rate of improvement for the single state model, the fast-learning state, and the slow-learning state separately for each of the four conditions.

**Fig 4.**
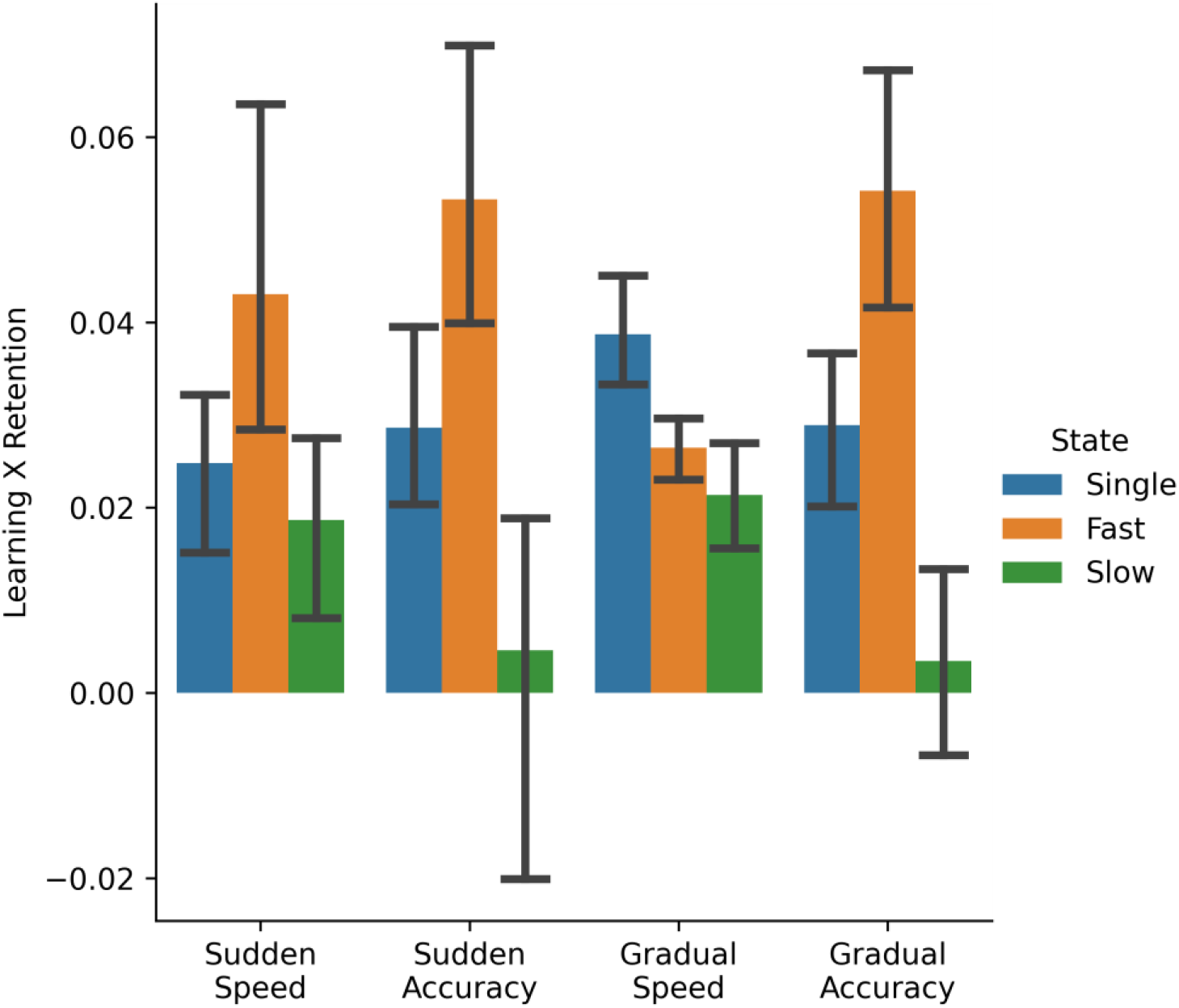
The multiplication of the best-fitting learning rate (B) and retention rate (A) parameter values, as a proxy for the rate of accuracy improvement, for each of the three states (the single state, the fast-learning state, and the slow-learning state). As seen in the figure, only for the gradual speed condition does the fast-learning state have a lower rate of improvement than the single state. In this case, the fast- and slow-learning states add up to approximately the same rate of improvement as the single state, as expected in a situation that does not require two states. Furthermore, the rate of the improvement for the fast-learning state in the gradual speed condition is similar to that of the slow-learning state for the other conditions, as expected if the gradual speed condition involves only slow, implicit learning.

As Figure 4 shows, the rate of improvement for the three states takes on a qualitatively different pattern for the gradual speed condition. For the other three conditions, the fast-learning state has a higher rate of improvement than that of the single state model (*t*(14) = 4.27,*p* < 0.001). In contrast, for the gradual speed condition, the rate of improvement for the fast-learning state is significantly *lower* than that of the single state (*t*(44) = 3.37,*p* = 0.002). Furthermore, in the gradual speed condition, the rate of improvement for the fast- and slow-learning states are approximately equal and approximately half the value of the rate of improvement for the single state model. This is expected in a situation, described earlier, where the dual state model mimics the single state model (e.g., when B_f_ + B_s_ = B and A_f_ = A_s_ = A). This corroborates the model comparison results that favor the single state in this condition. More importantly, the rate of improvement for the fast-learning state in the gradual speed condition is comparable to the rate of improvement of the slow-learning state from the other three conditions. In other words, the learning curve in the gradual speed condition requires only a single state with a slow rate of learning, similar to the slow, implicit learning state of the other conditions.

## Experiment 2: Removal of Online Visual Feedback During Motion

Experiment 1 identified circumstances that produced implicit learning without explicit learning (the gradual speed condition). Continuous visual feedback was provided during motion based on prior work finding that this leads to a larger contribution of implicit learning [10, 14, 54]. In the case of the gradual introduction of rotation, this continuous visual feedback may be crucial for blocking awareness of each additional 10° rotation. In other words, because participants can see when their motion first deviates from the desired direction, they may automatically adjust as they go, and fail to realize that the mapping between the track pad and onscreen direction was modified. However, if there is no online visual feedback during motion, participants will not adjust midstream and will simply miss the target by a full 10°. Because these 10° errors will occur in a systematic direction, the rotation may become noticeable even with speed emphasis, allowing the explicit system to contribute to learning. To test this hypothesis, Experiment 2 was identical to Experiment 1 except that visual feedback was removed during motion. Instead, participants were given only final outcome feedback after their motion was completed. If the removal of online feedback elicits explicit learning, the gradual speed condition should become similar to the other conditions. All experimental methods were the same as Experiment 1, except for the removal of online visual feedback, and other minor differences as noted. Because participants did not get continuous feedback, they made straight “shooting” movements to hit the target. Thus, movement error was computed as the angle formed by the lines joining the start point and the target and the line joining the start point and the actual end point. All modeling procedures were the same as for Experiment 1.

### Methods

#### Participants

Sixty-four individuals participated, with 16 in each of the four conditions.

#### Paradigm

The experimental task was built using the Psychopy platform [53]. On each trial, participants were shown a circular band formed by two concentric white circles around the center of the screen. A trial started with the appearance of a red circular cursor at the center of the screen along with a red circular target within the white-edged band. Participants were asked to make ballistic “shooting” movements to hit the target using the stylus. Once the movement was initiated, the dot at the center of the screen disappeared. Next, once the heretofore invisible movement crossed the inner boundary of the circular band, the dot reappeared at the corresponding location between the inner and outer circular bands, providing visual end-point feedback The target remained on the screen for the entire duration of the trial.

### Results

#### Performance after Learning

As seen in Figure 5, there was a main effect of the speed/accuracy emphasis manipulation after the 9 blocks of rotation learning, with smaller angular errors for the accuracy groups (*F* = 8.6,*p* = 0.004). However, in the subsequent washout block in which the rotation was removed, there were no apparent differences between the conditions. Unlike prior results [7], and unlike Experiment 1, the participants who received a gradual introduction of the rotation did not make larger errors than those who learned the rotation suddenly (*F* = 0.07, *p* = 0.78). This failure to find a larger washout effect for the gradual conditions is expected if the elimination of visual feedback during motion resulted in explicit learning for all conditions.

**Fig 5.**
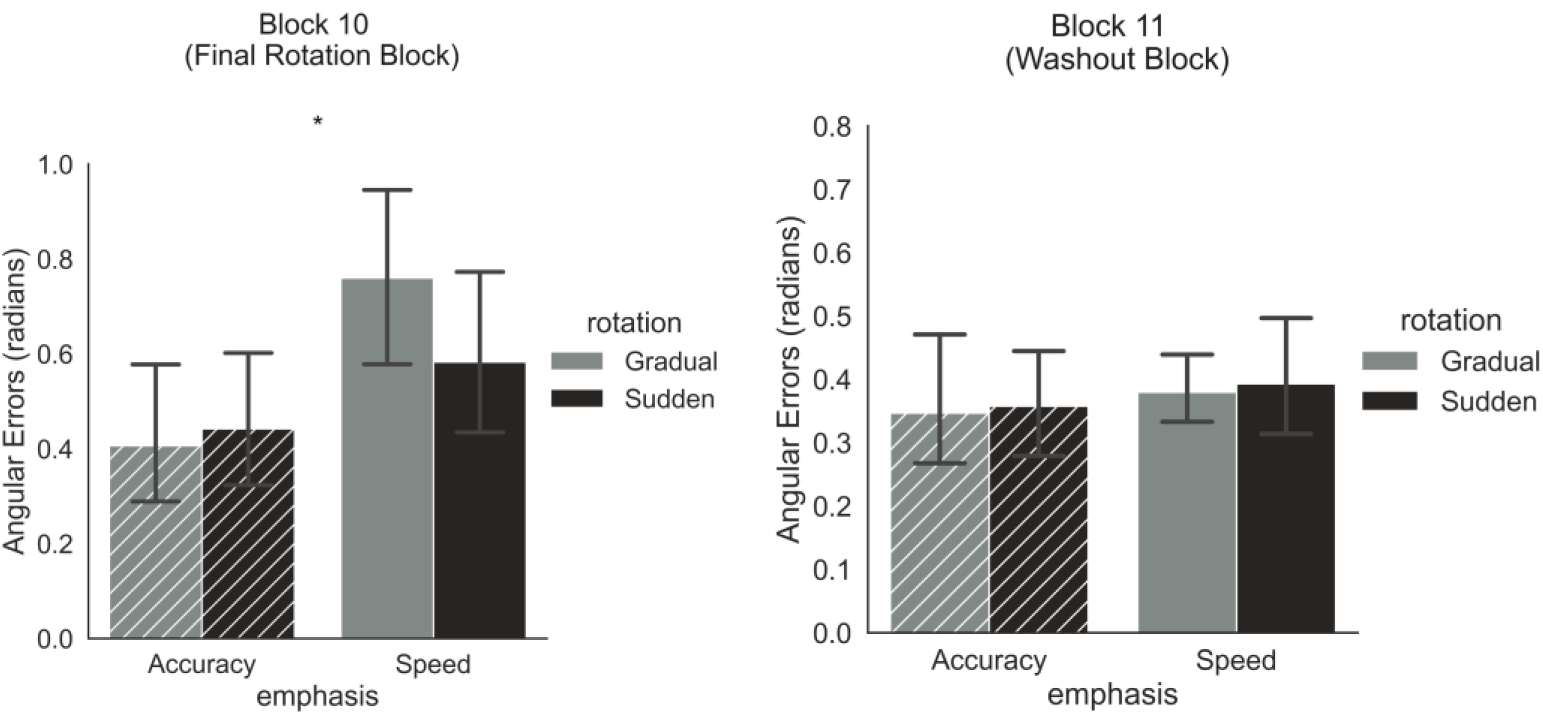
Accuracy after learning in Experiment 2. Left Panel: During the final block of rotation, participants in the speed conditions made larger errors than participants in the accuracy conditions for both the gradual and sudden introduction of rotation, demonstrating that the speed/ accuracy instructions affected behavior as expected. Right Panel: Unlike Experiment 1, there were no apparent differences between conditions during the washout block.

#### Model Comparison of Learning Curves

Each model was applied to the trial-by-trial rotation learning data shown in Figure 6, using the same fitting routines and model comparison metrics as Experiment 1.

**Fig 6.**
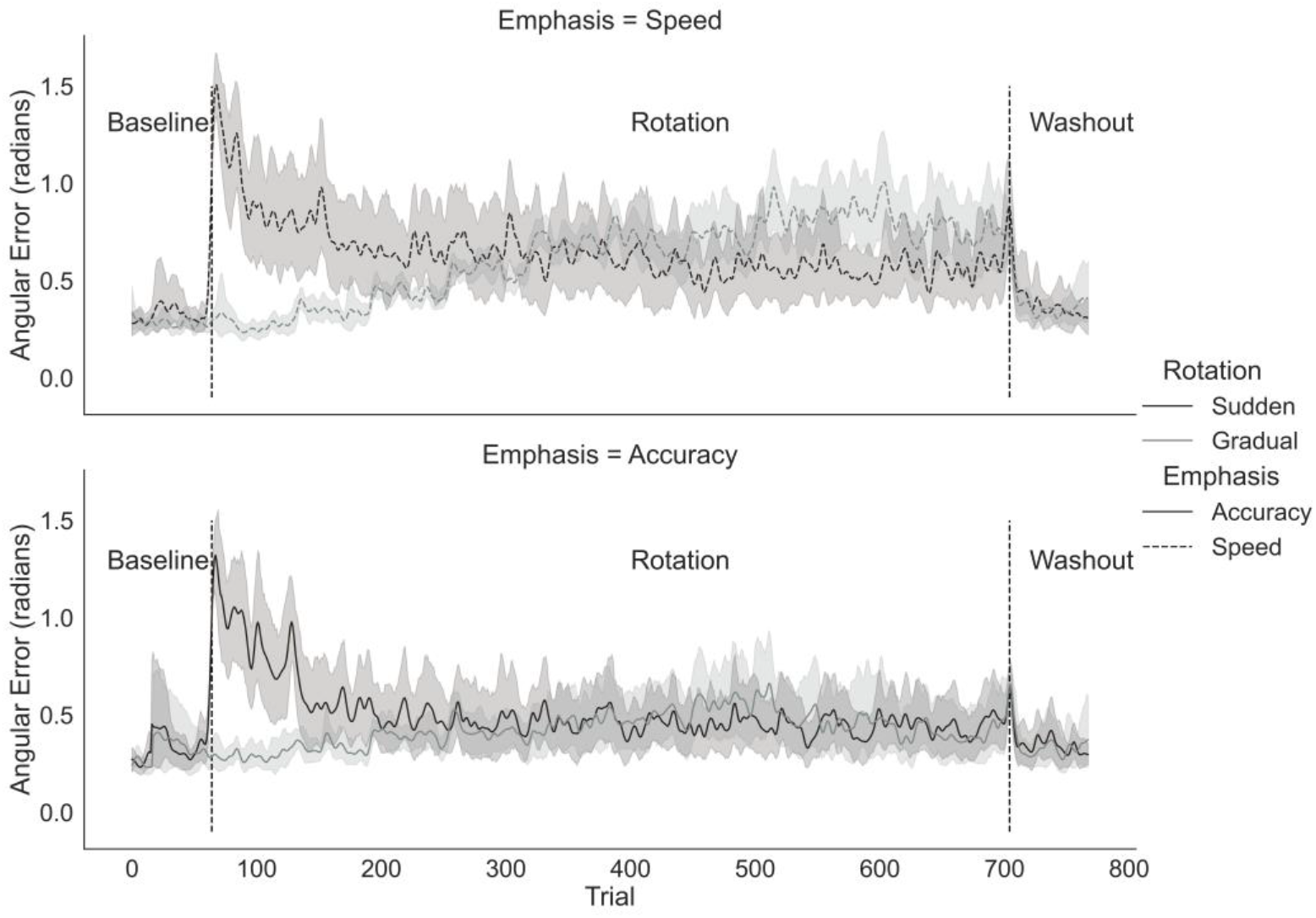
Average absolute angular motion direction error for the four groups of participants as a function of trial for Experiment 2. Darker lines indicate the groups that experienced a sudden introduction of the 90° rotation, and lighter lines the groups that were introduced to the rotation gradually. Top Panel: Results for speed emphasis groups (dotted lines). Bottom panel: Results for accuracy emphasis groups (solid lines). Shaded regions indicate the 95% confidence interval

As reported in Table 2, there were no apparent differences between the different conditions in terms of which model was preferred. More specifically, when using chi-squared, AIC, or BIC, the single state model was almost always preferred, regardless of condition. In contrast, when using cross-validation, more than half of the participants in each condition were better explained by the dual state model. As outlined above, the parameter-counting metrics (AIC, BIC and chi-squared) may unfairly penalize the dual-state model, whereas the cross-validation metric is likely to appropriately take account of true model flexibility. The cross-validation results might therefore be considered more reliable in this case of conflicting outcomes, suggesting that two states are operating in many participants. Most importantly, however, it is no longer the case that the gradual speed condition is different from the other conditions, in terms of model comparison.

**Table 2.**
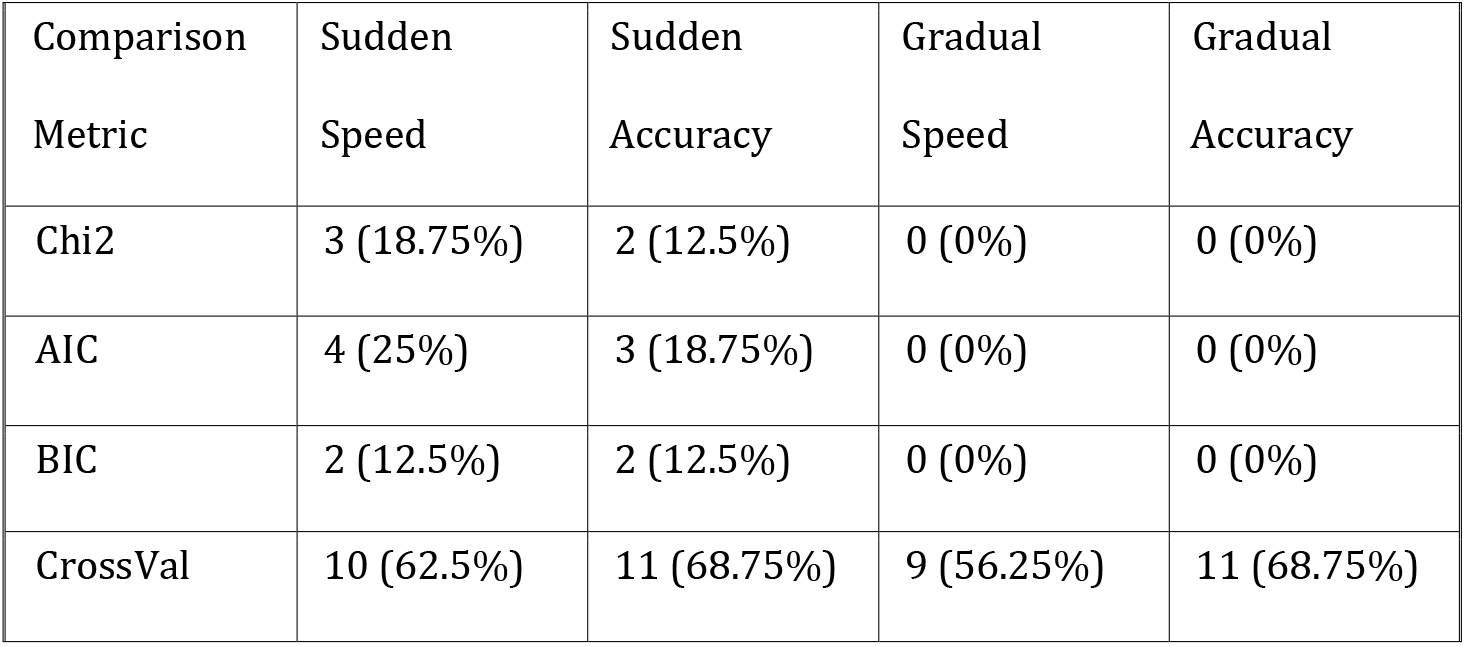
Model Comparison Results for Experiment 2 Showing How Often the Dual State Model was Preferred

As seen in Figure 7, the rate of improvement measure (given by the learning parameter multiplied by the retention parameter) shows a very similar pattern across all conditions, especially regarding the comparison of the single state and fast-learning state parameters. That is, in all conditions, there is no apparent difference between the rate of improvement for the single state model and the fast-learning state of the dual state model. Moreover, the rate of improvement for the slow-learning state is generally close to zero, whereas the fast-learning state’s rate of improvement mimics that of the single state model. As discussed above, one way in which the dual state model can mimic the single state model is when the slow-learning state is completely switched off, such as by setting either learning to zero or retention to zero. However, the (cross-validation) model comparison results indicate that two states are needed in a substantial number of participants. Thus, the best explanation for these results seems to be that both states are needed, and that learning in this paradigm is primarily accomplished by the fast, explicit state, but with a small amount of slower, implicit learning in some or all participants. This dominance of the fast-learning state likely explains the equivalence of the rate of improvement for the single state model and the fast-learning state of the dual state model (Figure 7).

**Fig 7.**
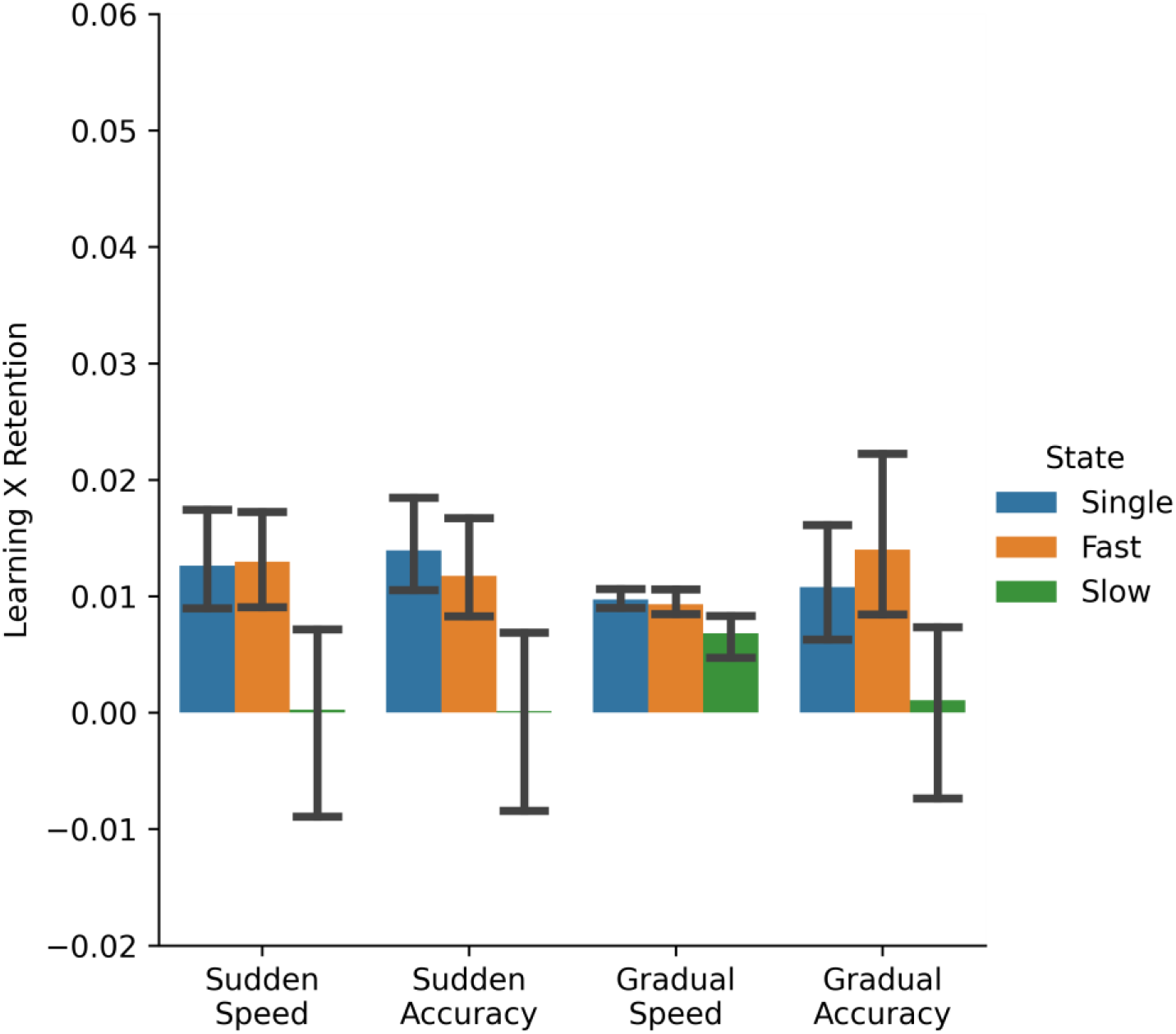
The multiplication of the best-fitting learning rate and retention rate parameter values – a proxy for the rate of accuracy improvement – for each of the three states (the single state, the fast-learning state, and the slow-learning state) in Experiment 2. All conditions show similar patterns wherein the rate of improvement for the single state model is very similar to the fast-learning state of the dual state model. The slow-learning state contributes minimally, or less than the fast-learning state, to the overall improvement, suggesting that learning is mostly through the fast-learning state when continuous feedback is not provided.

Finally, we note that the rate of improvement metric in Experiment 2 shows substantially lower values overall than in Experiment 1. This is not surprising considering that in the absence of movement feedback, Experiment 2 is much more of a guessing game for participants as they determine which direction to move to hit the target (errors were generally much higher and more variable). Without visual feedback on the position of the cursor during the hand movement, the opportunity for learning of either kind is much reduced. In Experiment 1, with movement feedback, not only was there an opportunity for the implicit system to experience the mapping between vision and motion in the course of the movement, but strategic midstream corrections during movement, made possible by the feedback, likely provided a more robust error signal for explicit learning, as well.

### Experiment 2 Discussion

This follow up study investigated the impact of continuous feedback on the contribution of the fast-learning state of the dual state model. We assessed the magnitude of the difference in aftereffects between the sudden and the gradual group of participants and looked for differences in model fits between the four conditions. We found that when feedback was not continuously available to the participants, there was no difference in how often the dual state model was selected between the experimental conditions. Particularly, participants in the gradual speed condition no longer showed a significantly lower dual-state model selection than the other conditions, as they did in the continuous feedback version of Experiment 1.

A possible explanation of these results is that the lack of continuous feedback removes visual information that would otherwise interfere with acquiring explicit knowledge about the experimentally induced rotation in the environment. That is, even with small rotations of 10°, as were provided in the gradual conditions, participants in the no-feedback paradigm might be better able to notice that the end-point feedback they receive is consistently divergent from the movement they perform, than in the paradigm with feedback (Experiment 1). This may have amplified the ability of Experiment 2 participants to explicitly ascertain the amount of counter aiming they needed to do to successfully hit the target. Such an explicit aiming strategy would influence the interactive dual state model and allow for the fast, explicit component to make a greater contribution to the final estimate of rotation than under conditions of continuous feedback.

## General Discussion

Prior studies found that visuomotor adaptation occurs automatically to a change (e.g., a rotation) in the mapping between the required motor direction and the onscreen position of visual targets in a reaching task [7,10,19,23]. Such implicit learning appears to be obligatory [38,39] and slowly-learned [3,23], resulting in slow extinction of the newly learned visuomotor mapping when it is subsequently removed. In contrast, explicit learning can produce accurate motions after just a few trials and can be quickly modified when the rotation is eliminated [3,23]. Thus, theories that include both slow learning and fast learning on every trial provide a good explanation of visuomotor adaptation behavior [3,9]. However, it was not previously known whether explicit visuomotor adaptation is similarly obligatory. Are there situations in which visuomotor adaptation is purely implicit, or does explicit learning always contribute? To determine whether the contribution of the fast explicit system can be reduced or eliminated, we tested participants’ adaptation to a 90° rotation under a range of conditions, defined by how the rotation was introduced (suddenly versus gradually) and by the instructions for how to optimize performance (moving quickly versus accurately). We found that the combination of gradual rotation learning with speed emphasis produced learning curves that were best explained by a single, slow-to-learn system, in the absence of any fast, explicit learning. Thus, when participants were encouraged to “leap before you look”, learning appeared to only reflect the slow, implicit system. This slow learning was also slow-to-forget, as indicated by a larger “aftereffect” in the washout phase in which the rotation was abruptly removed. We infer from this that, unlike the obligatory slow-to-learn, implicit system, the contribution of the fast-to-learn system is flexible and can be up-regulated or down-regulated with the right combination of experimental variables.

A variety of experimental variables have been examined in visuomotor adaptation reaching tasks. For example, the degree of rotation to which participants adapt during learning varies across different studies, from a small rotation of 8° [10], to a medium rotation of 20-60°, [12–14,20,23,40–45], to a large rotation of 75-90° [7,38,39,46]. Indeed, it has been shown that explicit learning scales with degree of rotation, whereas implicit learning does not [12]. Large rotations can be introduced suddenly or they can be introduced gradually [9,28,47–49], and a gradual introduction produces a larger aftereffect (i.e., larger errors in direction opposite to rotation) in a washout phase in which the rotation is removed [7]. In some studies, participants were given online visual feedback of their motions [7] while in others they were given only feedback about the final position of their motion [23]. Finally, some studies required that participants wait after the target appears before starting to move [15,26]. However, none of these studies asked participants to prioritize the speed of their motions. Because our study sought to maximize implicit learning, we attempted to minimize explicit awareness and time to explicitly plan motions by using a gradual introduction of the rotation, continuous visual feedback, and instructions to move quickly without delay. To replicate prior studies that found evidence for the necessity of both a fast-learning, explicit system and a slow-learning, implicit system, we also included conditions that we expected to increase explicit learning; namely, emphasizing accuracy over speed, and introducing the rotation suddenly to promote awareness of the rotation. Our results indicate that speed emphasis instructions reduce explicit learning, but only under conditions that also reduce explicit awareness of the rotation, such as by introducing the rotation gradually.

To determine whether the fast-to-learn explicit system was needed to explain behavior, we used formal model comparison of learning curves, comparing a single state model to a dual state model that included both a fast- and slow-learning system that combined to produced behavior. The dual state model is generally well-accepted and explains many visuomotor adaptation results. For instance, the dual state model has been found to offer a better explanation than the single state model of de-adaptation when the rotation is removed (washout) and of interference/transfer effects when participants were asked to subsequently learn a different rotation after adaptation [3]. Because the single state model is a special case of the dual state model, the dual state model is inherently more flexible (i.e., it can explain a wider variety of behaviors) and model comparison requires some form of penalization for this extra flexibility, to determine if both states of the dual state model are needed. Four different methods of model comparison were used, applied separately to each participant. In Experiment 1, all four methods indicated that the learning curves of the participants assigned to the gradual speed condition were well described by the single state model. In this case, the goodness of fit improvement from adding a second learning state was negligible. Thus, parsimony supports the conclusion that the fast-learning state did not contribute to behavior in this condition. In contrast, nearly all of the participants in the sudden accuracy group produced learning curves that were indicative of both fast (e.g., a rapid drop in accuracy over a few trials) and slow (e.g., a very slow additional improvement over hundreds of trials) learning, and thus better explained by the dual state model. The other two conditions (sudden speed and gradual accuracy) reflected a more even mixture, with some participants better explained by the single state model and others by the dual state model.

Our finding that the single state model provides a sufficient explanation in some situations does not necessarily specify whether that single learning state is implicit (slow) or explicit (fast). However, an analysis of the best-fitting parameter values can address this question by comparing the rate of improvement (approximated by the product of the learning and retention parameters of the model) for the fast-learning state of the dual state model to the rate of improvement for the single state model. This comparison is instructive because, in cases where the dual-state model is mimicking the single state by setting one state to zero, all learning will load onto the “fast-learning state” parameters (owing to Eq.7). In Experiment 1, the rate of improvement for the fast-learning state was greater than that of the single state model for all three conditions except the gradual speed condition. In contrast, in the gradual speed condition, the fast-learning state had a lower rate of improvement than the single state model, and the fast- and slow-learning rates of improvement together summed to approximately the rate of the single state model, suggesting that the dual state model was mimicking the single state model, Furthermore, the fast-learning state’s rate of improvement in the gradual speed condition was comparable to that of the slow-learning state in the other three conditions. Thus, the learning in the gradual speed condition was consistent with a single system of slow, implicit learning.

### Model-Based and Model-Free Learning

Implicit, slow learning versus explicit, fast learning in visuomotor adaptation are closely related to different kinds of computational reinforcement learning models termed “model-free” versus “model-based”, respectively [55,50]. Put simply, model-free is trial-and-error learning-by-doing, whereas model-based involves mental simulation of possible outcomes before choosing between action alternatives. As might be expected, model-free learning is very slow and inflexible whereas model-based learning more quickly adapts to changes in the environment. Furthermore, because model-based learning requires some form of mental simulation to calculate the subsequent outcomes that may arise from different actions (e.g., different movement directions), it follows that an emphasis on speedy action rather than accuracy may promote greater use of model-free learning. Our results mesh with this literature, suggesting that a model-free system that slowly learns inflexible representations can dominate the learning of a gradual motor adaptation under conditions of speed emphasis.

Analogous to the dual-state model, it has recently been proposed that model-free and model-based learning can operate in a coordinated fashion, perhaps with model-based learning serving as the offline teacher of the model-free system [51,52]. When applying the dual state model to learning curves that appear to require both states, the fast-learning state rapidly learns the rotation, explaining the initial rapid reduction in error. But after many learning trials, the slow-learning state eventually learns the rotation and is able produce fast accurate movements without any help from the fast-learning state. Once this happens, the fast-learning state loses its knowledge of the rotation because there is no longer enough movement error to maintain its estimate of rotation considering that the fast state is also fast to forget the rotation. Thus, there is a transition of knowledge regarding the rotation from the fast- to the slow-learning state. This transition of knowledge is similar to the proposal that the model-based system can serve as the teacher of the model-free system. Our results indicate that the fast-learning, explicit system is not always needed to explain behavior. By analogy, this suggest that some learning situations (e.g., a need for fast decisions during gradual change of the environment) may short-circuit the explicit, model-based learning. However, even when model-based learning unavailable, the model-free system continues to learn through direct experience.

## Conclusions

In this article, we explored the role of implicit and explicit learning in adaptation to novel motor mappings. While standard models of motor learning suggest that humans need both implicit and explicit components to successfully adapt to new motor mappings, we show that when the new motor mappings are introduced gradually, under time pressure and with online feedback, participants may be able to adapt without using explicit learning. We conducted two behavioral experiments: first, participants learned a visuomotor adaptation task under varying task emphasis instructions and varying perturbation schedules, along with continuous visual feedback on their movements; second, participants performed the same task under the same conditions, without the continuous, online visual feedback.

Replicating prior results, we found that in the continuous feedback case participants who adapt to the rotation gradually under time pressure find it harder to go back to veridical reaching. Our model comparison results suggest that these participants rely on a single system – a slow-learning, implicit system – more than the groups that learned under other conditions. We do not find similar systematic differences in participants’ reliance on two states when continuous feedback is not available, which indicates the importance of continuous feedback for engaging implicit learning (or for interfering with explicit learning).

## Acknowledgements

This work was supported by NIH grant (RF1MH114277) awarded to PIs Cowell and Huber. We would also like to thank all the undergraduate Research Assistants of the cEMNL-cMAP lab in their help in data collection.

